# Association between use of the progestin Depot Medroxyprogesterone Acetate and Cervicovaginal Lavage Proinflammatory Cytokines among HIV-1 and HSV-2 co-infected African women: a case control study

**DOI:** 10.1101/045344

**Authors:** Edwin Walong, Christopher Gontier, Walter Jaoko, Elizabeth Bukusi

## Abstract

**Background:** Use of the progestin contraceptive Depot Medroxyprogesterone Acetate (DMPA) by HIV 1 infected women is associated with increased female to male transmission of HIV. Mucosal innate immune activation has been proposed as a likely mechanism. To establish the effect of DMPA upon mucosal immune activation, this study sought to evaluate the concentrations of 5 proinflammatory and the regulatory cytokine IL 10 in cervicovaginal lavage fluid.

**Methods:** This was a case control study, 70 participants were recruited, comprising of 35 asymptomatic ART naïve HIV positive women on DMPA recruited as cases and 35 age matched asymptomatic ART naïve HIV positive women not on contraceptives recruited as controls. Peripheral blood CD4 and total lymphocyte counts, High vaginal swab microscopy, endocervical smears and cervical cytology were performed for each participant. Concentrations of six proinflammatory cytokines were measured on cervicovaginal lavage by multiplex cytometric bead array.

**Results:** The mean age of cases was 26.8 years and 30 years for controls. Total lymphocyte counts and CD4 cell counts were significantly higher among cases (p=0.02 and 0.004 respectively). HSV 2 prevalence as determined by ELISA was higher (p=0.034 among cases. The concentrations of the cytokines IL 1β, IL 6, IL 8, IL 12p70 and TNF α were lower among cases, with IL 1β being statistically significant (p=0.046). Concentrations of IL 10 was higher among cases (p=0.022). On multivariate analysis, reduction in IL 1β and IL 8 were associated with the duration of DMPA use (p=0.015 and 0.041 respectively). Inclusion of HSV 2 into the multivariate models showed elevation of all cytokines measured (p=<0.001).

**Conclusion:** DMPA use is associated with reduction of proinflammatory cytokines and elevation of the regulatory cytokine IL 10. This may explain increased female to male transmission of HIV infection by modulation of male genital tract mucosa in the absence of increased HIV 1 genital shedding.

## BACKGROUND

The HIV pandemic in sub-Saharan Africa disproportionately affects women who constitute over 60% of persons leaving with AIDS in Africa (1). The progestin Depot Medroxy-Progesterone Acetate (DMPA), a widely used contraceptive in Eastern and Southern Africa, is associated with a higher rate of female to male transmission of HIV, presenting a potential risk of new infections among male partners of infected women (2). The mechanisms for increased female to male transmission of HIV, particularly those due to alterations in mucosal immune activation within the female lower genital tract are poorly understood (3). Determination of this mechanisms are important for evaluation of safety of hormonal agents as well as their effect on susceptibility to HIV and other infections in which mucosal immunity provides the initial barrier to infection.

Innate mucosal immune activation is the likely mechanism of increase in to HIV 1 and HSV 2 (4). Studies have shown an increase in lower female genital tract mucosal immune cells suggesting mucosal immune activation and elevation of proinflammatory cytokines (5). Among women living with HIV, epidemiological studies have shown higher odds of female to male transmission of HIV, which is not associated with increased systemic or genital viral loads (2). Studies in animal models and cell lines show DMPA suppresses the synthesis of type 1 and type 2 cytokines (6) (7). In this study, we report the concentrations of proinflammatory, specifically IL 1*β*, IL 6, IL 8, IL 12 and TNF α, as well as IL 10, a regulatory cytokine among women living with HIV using DMPA.

## METHODS

This was a case-control study conducted at the Family Aids Care and Education Services clinic, an outpatient comprehensive care clinic in Kisumu, Kenya. Study participants were HIV positive Anti-Retroviral Therapy naïve women aged between 18-45 years, World Health Organisation (WHO) stage 1 or 2, not on active treatment regimens for opportunistic infections, whose peripheral blood CD 4 counts were above 350,000 per litre. Cases were women in whom DMPA had been administered through deep intramuscular injection within three months prior to recruitment. Controls were women who were not using hormonal contraception.

Using a standard questionnaire, demographic and social information was obtained. Total Lymphocyte Counts and CD 4 lymphocyte counts, performed as part of comprehensive care, were obtained from electronic medical records. Physical examination was then performed, which included visual evaluation for evidence of ulcerative sexually transmitted infections.

Specimens collected included 4ml of whole blood obtained by venepuncture from the antecubital veins. After insertion of a sterile vaginal speculum, high vaginal swabs and endocervical swabs were obtained using a Dacron tipped swabs. The high vaginal swab was transported in sterile tubes and transported to the laboratory within 30 minutes for wet mount microscopy and Gram stains for diagnosis of *Trichomonas vaginalis, Candida sp* and Bacterial vaginosis. Endocervical swab samples were inoculated into Thayer-Martin Culture media for *Neisseria gonorrhoeae* culture. Cervico-vaginal lavage fluid (CVL) was obtained by introduction of 10ml of sterile phosphate buffered fluid into the vagina, allowed to pool in the cervix and vaginal fornices, and aspirated using a pipette. Recovery of more than 8ml was considered successful. This was transported to the laboratory in sterile tubes for processing.

Laboratory diagnosis of Herpes simplex type II infection was by ELISA using antibodies against HSV 2 specific g-glycoprotein (Kalon™ Diagnostics) Diagnosis of Bacterial Vaginosis was by microscopy of Gram stained direct High Vaginal Smears using the Nugent Criteria. Wet mount preparations and microscopy of HVS was performed for diagnosis of infection by *Trichomonas vaginalis* and *Candida spp*. The CVL was centrifuged at −4 degrees Celsius, separated and divided into 1 ml aliquots, seven bearing the supernatant and one cell pellet. These were frozen at −80 degrees Celsius.

For determination of the cytokines IL 1*β*, IL 6, IL 8, IL 10, IL 12 p 40 and TNF *α*, frozen CVL specimens were transported to the Kenya Aids Vaccine Initiative Laboratory, University of Nairobi, where they were thawed at room temperature. Using the Mulitplex Cytometric Bead Array (CBA) proinflammatory cytokine assay kit (BD Biosciences, Lot no 551811), incubation with the primary antibody coated beads was performed according to the manufacturer’s instructions. These were acquired using a Flow Activated Cell Sorter (FACS) Calibur ™ (BD Biosciences). These were evaluated using the BD Cellquest ™ software (BD Biosciences) and concentrations expressed in ng/ml.

Data was analysed using the Statistical Package for Social Sciences^®^ version 17. Bivariate statistics were performed to evaluate the objectives. These included student t test for normally distributed data and Mann-Whitney u test for skewed data. Multivariate multiple regression analysis was performed for association of HSV 2 latency and Cytokine concentrations.

Prior to commencement of the study, clearance was obtained from the Kenyatta National Hospital/University of Nairobi Scientific and Ethical Review Committee. Informed consent was obtained from each study participant. Confidentiality was maintained.

## RESULTS

### Demographic and Clinical Data

Demographic and Clinical data is presented in **table 1**. There were 35 cases and 35 controls. The prevalence of HSV 2 infection was 94% and 83% among cases and controls respectively. None of the participants screened had active HSV 2 lesions. Median CD 4 counts were 649 and 573 among cases and controls respectively.

**TABLE 1:** DEMOGRAPHIC CHARACTERISTICS OF CASES (WOMEN ON DMPA) AND CONTROLS.

|  | Variable | Cases (n=35) | Controls (n=35) |
| --- | --- | --- | --- |
| Age | Mean Age | 26.7 | 30.09 |
|  | Median Age | 26.0 | 28.0 |
|  | 95% C.I. for age | 24.8-28.9 | 27.6-32.5 |
|  | SD of Age | 5.9 | 7.04 |
| Marital Status | Married | 20 | 23 |
|  | Single | 8 | 7 |
|  | Separated | 3 | 4 |
|  | Widowed | 4 | 1 |
| Highest Educational Level Attained | Primary School | 20 | 23 |
|  | Secondary school | 10 | 11 |
|  | Middle level college | 4 | 1 |
|  | University | 1 | 0 |
| Occupation | Professional | 7 | 4 |
|  | Housewife | 2 | 4 |
|  | Farming | 2 | 1 |

### Cytokine Concentrations in CVL Supernatant

To detect the effect of DMPA use on mucosal immune activation, we determined the concentrations of cytokine concentrations among cases and controls (presented in **table 2**). Participants on DMPA had higher IL 10 concentration. The concentrations of IL 1β was reduced among cases, this was statistically significant. There was a non-statistically significant reduction in IL 6, IL 8, IL 12 and TNF α among cases. Multivariate analysis showed time dependent reduction in the concentrations of IL 1β and IL 8 (**Figure 1** **and** **Figure 2**).

**TABLE 2:**
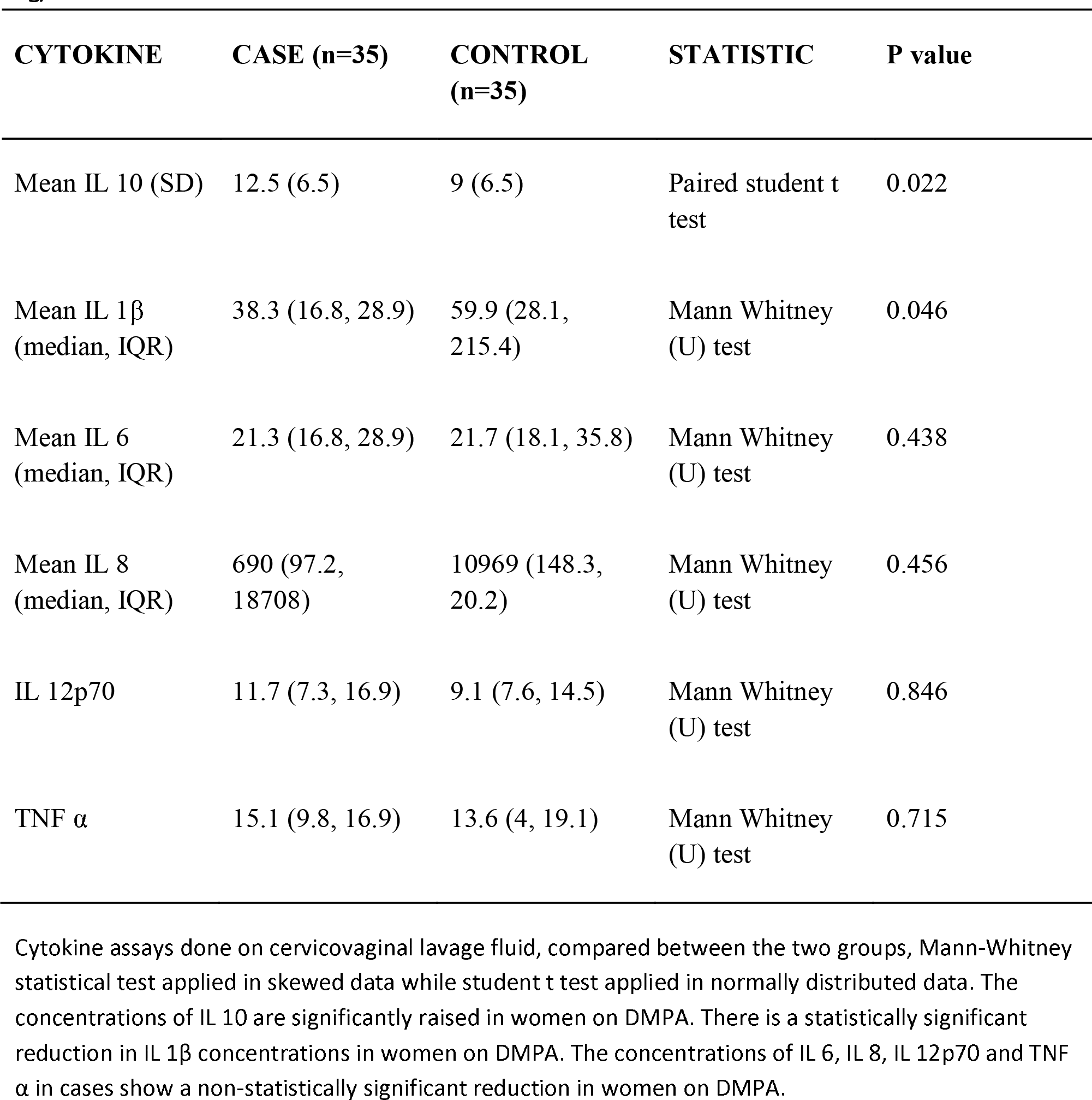
CONCENTRATIONS OF CERVICOVAGINAL LAVAGE FLUID PROINFLAMMATORY CYTOKINES IN ng/ml.

**FIGURE 1:**
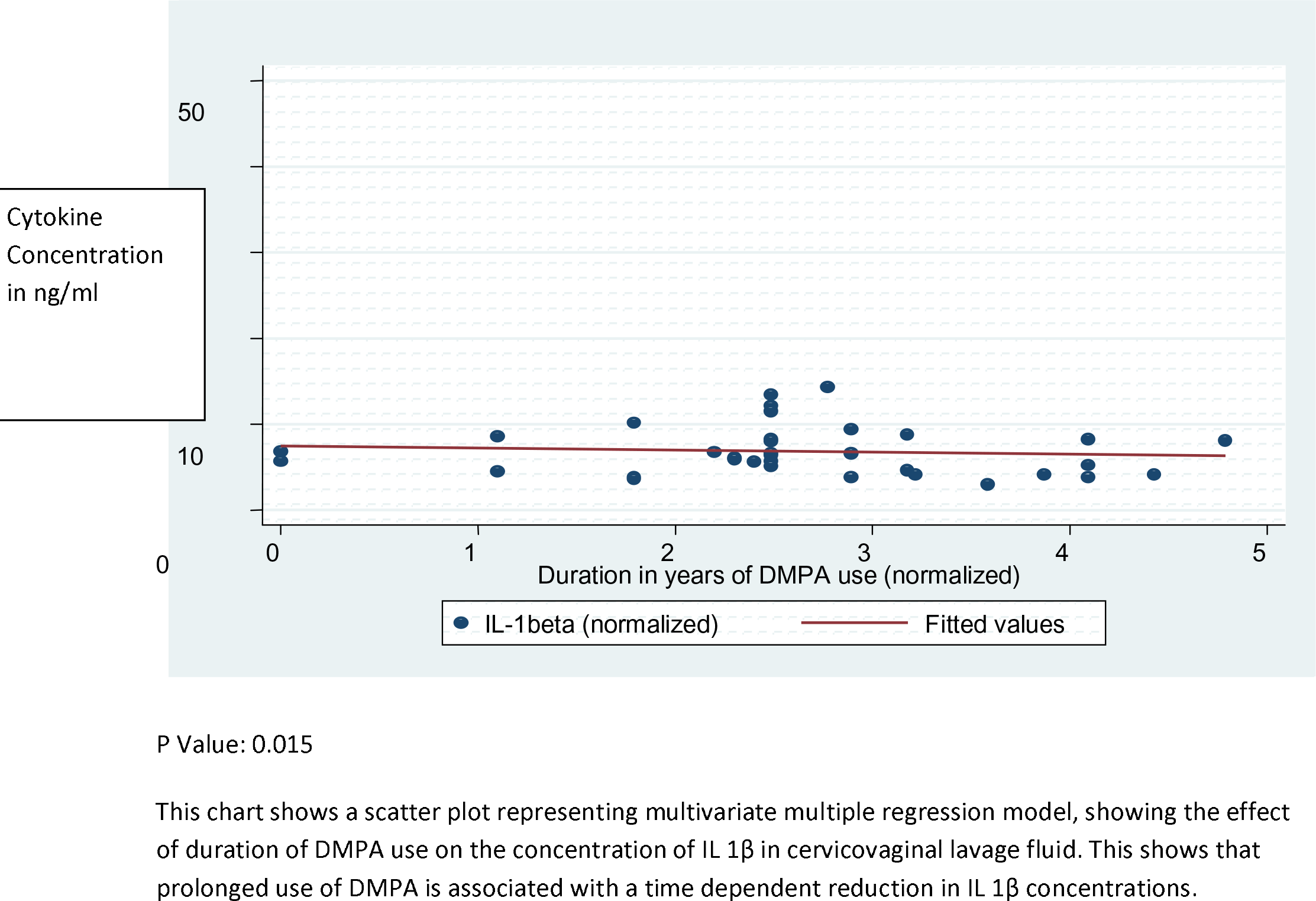
SCATTER PLOT SHOWING CORRELATION BETWEEN IL 1 BETA AND DURATION OF DMPA USE. P Value: 0.015

**FIGURE 2:**
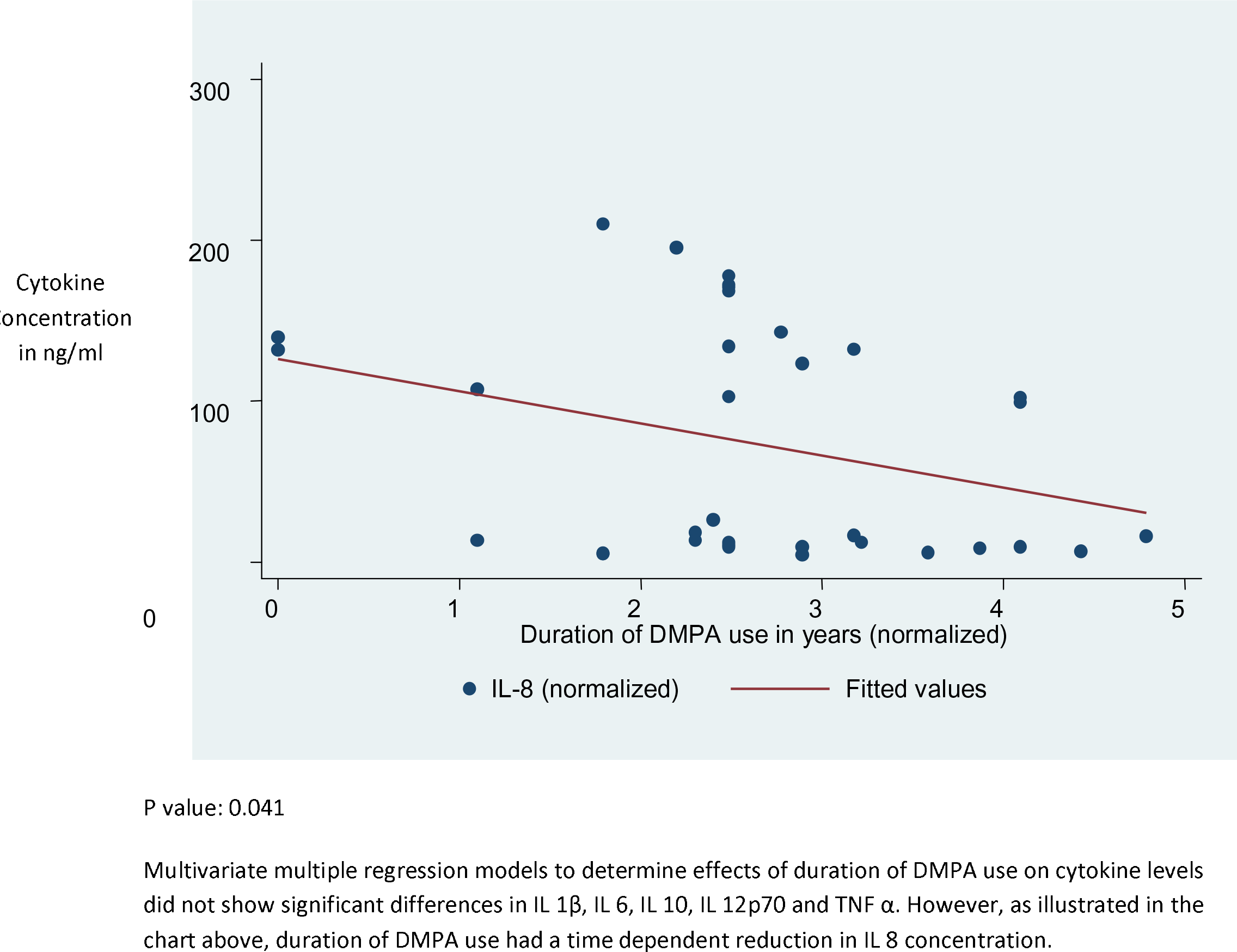
SCATTER PLOT SHOWING CORRELATION BETWEEN IL 8 CONCENTRATIONS (ng/ml) AND DURATION OF DMPA USE. P value: 0.041

### Effect of HSV 2 Co-Infection

To evaluate the effect of HSV 2 infection, Multivariate multiple regression analysis incorporating index values derived from HSV 2 ELISA was performed. This showed higher concentrations of all cytokines measured among HSV 2 infected women (**Table 3**).

**TABLE 3:** MULTIVARIATE MULTIPLE REGRESSION ANALYSIS OF THE EFFECT OF HSV 2 ELISA INDEX VALUES ON CVL CYTOKINE CONCENTRATIONS.

| **CYTOKINE** | **COEFFICIENT OF HSV 2 IgG** | **95% Confidence Interval** | **P value** |
| --- | --- | --- | --- |
| TNF $\alpha$ | 20.29 | 12.9-27.7 | <0.001 |
| IL 1β | 11.44 | 7.7-15.2 | <0.001 |
| IL 6 | 46.1 | 21.6-70.7 | 0.001 |
| IL 8 | 198.5 | 84.4-312.6 | 0.002 |
| IL 10 | 15.6 | 8.8-22.3 | <0.001 |
| IL 12p70 | 3.4 | 1.7-5.2 | 0.001 |

Represented on these tables are multivariate regression models evaluating the effect of the HSV 2 index values as determined by ELISA on CVL cytokines. This shows a strong association between HSV 2 and the cytokines measured in cervicovaginal fluid and as such presents a significant confounding factor. A higher HSV 2 index value is associated with elevation of all cytokines measured.

## DISCUSSION

This study showed an elevation in CVL IL 10, a regulatory cytokine and reduction in the concentrations of the proinflammatory cytokines IL 1β, IL 6, IL 8 and IL 12 among women on DMPA. Studies have demonstrated a dose dependent DMPA induced reduction in proinflammatory and T helper 1 (Th1) cytokines as well as suppression of regulatory and Th2 cytokines (8).

Elevation of IL 10 was unexpected. Animal models and cell lines exposed to show consistent reduction in cellular IL 10 synthesis (7) (8). Although HSV 2 infection is associated with elevated IL 10, a synergistic effect may have been due to DMPA use (3). Unexpected effects of DMPA has been observed due to its selective modulation of the Glucocorticoid receptor, where it displays agonist and antagonist effects (9) (10) (7).

Although inconclusive, various epidemiologic studies and metanalysis have shown higher odds of female to male HIV transmission due to DMPA use (2) (11). Recent studies have shown that DMPA use is not associated with increased HIV shedding among women living with HIV (Waithera Githendu, Unpublished Observations). The mechanisms of increased female to male transmission of HIV are not currently known (2) (11). Our study shows differences in mucosal immune cytokines such as DMPA induced elevation of IL 10 may provide the mechanism for modulation of male genital mucosal immunity. The functions of IL 10, a regulatory cytokine, include suppression of T helper lymphocyte, myeloid and dendritic cell functions resulting in increased susceptibility to infection. (7) (12).

Although we hypothesize that exposure of the penis IL 10 and HSV 2 are plausible mechanisms in DMPA associated female to male transmission of HIV, evidence of this is scant. Intra and inter individual variation in pharmacokinetics of DMPA after intramuscular injection of DMPA is a significant confounding factor (7). We showed trends of cytokine alteration due to prolonged DMPA use which may be consistent with prolonged exposure and effect. A longitudinal study would evaluate the likely mechanisms of DMPA alteration in mucosal milleau within the female genital tract and penile skin or mucosa of their male sexual partners.

## Author Contributions

EW was the principal investigator, developed the concept, designed the study and performed cytological evaluation cervical smears of cytometric bead array analysis of cytokines. WA performed client recruitment and counseling, microbiologic analysis of cervical specimens. CS participated in the design of the study and cytokine analysis. WJ participated in the design of the study, data interpretation and the writing of the manuscript. EB participated in the design of the study, conceptual framework, interpretation of data and the eventual writing of this manuscript. All authors read and approve of this manuscript.

## Acknowledgements

Alrica Atieno, May Maloba, Cirilus Ogola, Megan Huchko, Patrick Oyaro, Kevin Owuor for their assistance in recruitment and statistical support (Family Aids Care and Education Services, Research Care and Training Program, Kenya Medical Research Institute); Onyango Obila, Bashir Deko, Omu Anzala who provided bench space for cytokine analysis (Kenya Aids Vaccine Initiative, University of Nairobi), June Odoyo, Robert Bailey who provided logistical and laboratory support for the microbiologic studies (UNIM Kenya) Emily Rogena, Joshua Nyagol, Jessie Githanga (Department of Human Pathology, University of Nairobi). This study was supported by the Research Care and Training Program of the Kenya Medical Research Institute. The preparation of this manuscript was supported by Partnership for Innovative Medical Education in Kenya (PRIME-K/MEPI), NIH Grant Number 5R24 TW008889-02.

